# Subcellular Characterization of the Molecular Determinants of Ebola VP40 Trafficking and Assembly

**DOI:** 10.64898/2026.02.13.705837

**Authors:** Tyler Huth, Ella Wiggenhorn, Susmita Khanal, William Wan

## Abstract

Ebola virus is a single-stranded negative-sense RNA virus that can cause severe hemorrhagic fevers in humans. Ebola virus, along with other members of the filoviridae family, produce virions with a characteristic filamentous morphology. VP40, the filovirus matrix protein, is responsible for curving the host plasma membrane (PM). Expression of VP40 and the assembly of the matrix layer results in the budding of filamentous particles. VP40 forms cytosolic homodimers via interactions in its N-terminal domain, while interactions in its C-terminal domain drive oligomerization into the 2D-crystalline matrix layer. While VP40 is expressed throughout the host cytosol and assembles on the inner leaflet of the PM, VP40 does not appear to directly bind the PM but instead requires interactions with components of the host secretory machinery. Here, we characterize a series of VP40 mutants targeted to the molecular determinants of Ebola VP40 assembly and traWicking using confocal microscopy and genetically-encoded fluorescent tags. Using this approach, we characterize the subcellular distribution of these mutants, showing novel cellular phenotypes. Several mutants previously characterized as traWicking deficient show aggregation dependent on membrane binding, suggesting a possible route of VP40 traWicking. We then co-expressed these mutants with organelle markers, which provide insights into the aWected parts of the host traWicking pathways. Together, our results provide new insights into molecular interactions that drive the assembly of the Ebola matrix layer.

## Introduction

Ebola virus can cause severe hemorrhagic fevers in humans with case fatality rates ranging from 50% to 90% (1). Ebola virus is a member of the viral family filoviridae, which also includes other highly-lethal pathogens such as Marburg and Sudan viruses (2). Filoviruses possess single-stranded negative-sense RNA genomes, placing the family within the viral order Mononegavirales (3), which includes other important human pathogens such as measles and respiratory syncytial viruses.

Filoviruses take their name from the characteristic filamentous morphology of their virions. The formation of these filamentous virions is driven by the assembly of the viral matrix layer (Fig 1), a structured lattice composed of the viral matrix protein VP40 (4). Expression of VP40 is both necessary and suWicient to bud filaments from the plasma membrane (PM), indicating that the molecular determinants that drive homotypic assembly, as well as any host-pathogen interactions necessary for budding, are encoded in the structure of VP40 (5). VP40 consists of distinct N- and C-terminal domains (NTD and CTD, respectively) (Fig 1) (6, 7). VP40 is thought to primarily exist as a homodimer in solution (7), which is mediated by interactions between the NTDs (Fig 1B). VP40 CTDs contain a basic patch of 6 lysine residues (K221, K224, K225, K270, K274, and K275) which is thought to mediate association with the PM and are essential to matrix assembly and budding; residues K224 and K225 are in a disordered 10-residue loop within the patch (Fig 1C). As VP40 dimers accumulate at the inner leaflet of the PM, VP40 CTDs interact to form extended chains of dimers, which then stack to form 2D crystalline arrays (Fig 1A), i.e. the matrix layer (4). Assembly of VP40 into the matrix layer curves the PM, resulting in the formation of filamentous buds.

**Figure 1:**
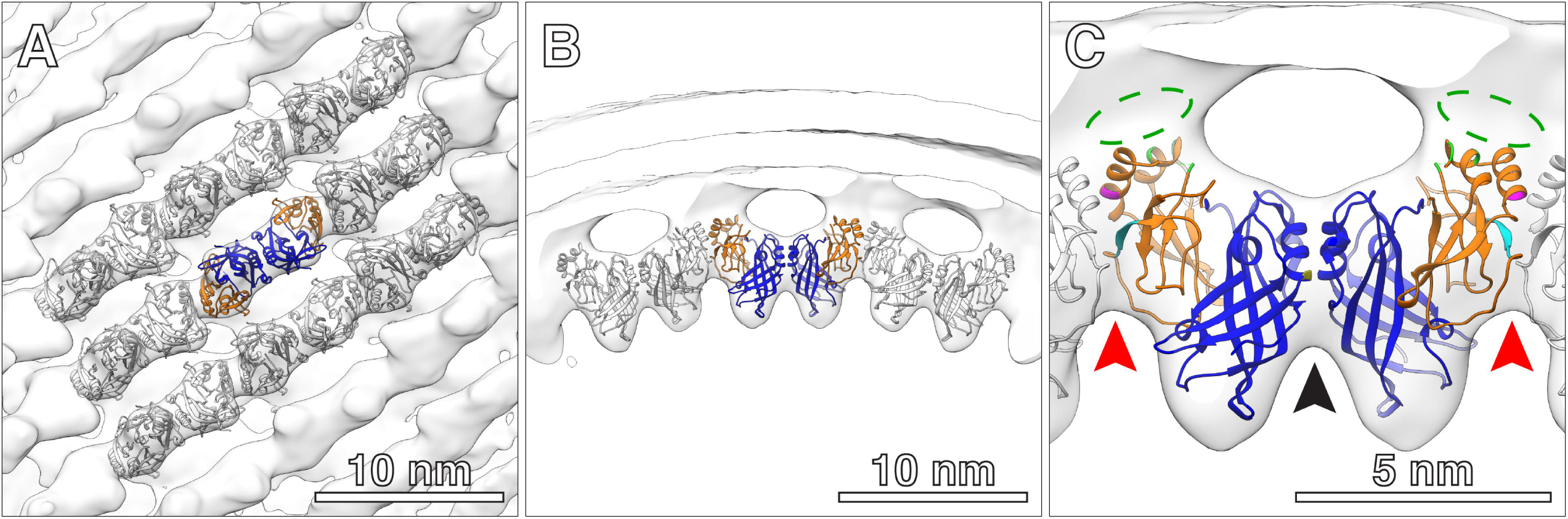
Structure of the Ebola Virus Matrix Layer. A) Cryo-electron tomography (cryo-ET) structure of the Ebola virus matrix layer within VP40 virus-like particles (VLPs). This view is from within the VLP and VP40 dimers are fitted into the cryo-ET density. Central dimer has NTDs colored blue and CTDs colored orange. B) View along a chain of VP40 dimers. The CTD densities show attachment to the inner leaflet of the VLP envelope. C) Detail of the VP40 heterodimer within the assembled matrix. Black arrowhead indicates the NTD dimerization interface and red arrowhead indicates the CTD oligomerization interface. Residue L117 is in yellow, R241 in magenta, residues 303 – 303 in cyan, and residues K221, K270, K274, and K275 in green. Residues K224 and K225 are within the unresolved loop, and are thought to be at the membrane-attachment point highlighted by the green ellipse. Cryo-ET maps are from Electron Microscopy Data Bank (EMDB) deposition EMD-11661 and molecular model is from the Protein Data Bank (PDB) deposition 4LDB.

In addition to its dimeric and matrix states, VP40 can also assemble into other isoforms. VP40 can form an RNA-binding octameric ring, which consists of the NTDs forming a structured central core, with the CTD’s flexibly “swung out” and unresolved in crystal structures (7, 8). RNA-binding is not necessary for matrix assembly, but appears to be involved in regulation of viral translation, making RNA-binding a requisite part of the viral life cycle (9). A linear hexamer form has also been characterized by X-ray crystallography; this isoform combines features of the dimeric and octameric isoforms to assemble a linear core of six NTDs, with one ordered CTD on each end and four “swung out” disordered CTDs in the center (7). It is unclear if the linear hexamer isoform serves a direct role in the viral life cycle.

As with all members of Mononegavirales, Ebola virus replication occur within cytosolic viral factories (10–13). Ebola viral factories are membrane-less molecular condensates, where nascent nucleocapsids assemble prior to traWicking towards budding sites at the PM (14, 15). Ribosomes appear to be excluded from viral factories, suggesting that viral mRNA is translated outside of viral factories (14). While VP40 ultimately assembles at the inner PM, it does not appear to bind the PM directly from the host cytosol (16). VP40 has been shown to directly bind Sec24c (16), a component of the COPII anterograde traWicking machinery that mediates transport between the endoplasmic reticulum (ER) and Golgi (17), as well as modulate the cellular distribution of Rab11 (18), a component of the endocytic recycling pathway. These interactions suggest that VP40 migrates to the PM by taking advantage of the host cell traWicking machinery.

In this manuscript, we characterize the subcellular distribution of VP40 mutants using confocal microscopy. Our VP40 constructs consist of super-folder green fluorescent protein (sfGFP) (19) fused to the N-terminus of VP40, which enables fluorescence imaging without the membrane permeabilization required in immunostaining approaches. The mutants we characterize were previously described in the literature as modifying the assembly of higher-order VP40 structures and VP40 interactions with Sec24c. Using our sfGFP-labelled VP40, we find a number of phenotypes not previously described in the literature, including several distinct subcellular aggregation patterns associated with Sec24c-binding mutations. Co-mutations to abrogate membrane binding precludes the formation of these aggregates, suggesting that aggregation occurs on membrane surfaces and may represent stalled traWicking intermediates. Co-expression of these mutants with markers for the ER, cis-Golgi, and trans-Golgi network showed co-localization or disruption of subcellular localization. Our results provide new insights into eWects of VP40 mutations and the molecular mechanisms by which VP40 is traWicked from the cytosol to the inner leaflet of the PM.

## Results

### Mutations at the Assembly Interfaces

Wild type (WT) sfGFP-labelled VP40 shows diWuse cytosolic signal with no distinct subcellular localization as well as intense signal localized to the budding filaments (Fig 2A). The high signal intensity of filaments is likely due to the very high density of sfGFP molecules in the ∼60 nm diameter filaments (5, 4). Depending on the orientation of filaments, some can take on a “beads-on-a-string” appearance in our images, which is due to finite Z-slicing of thin filaments during confocal microscopy data collection. Western blotting confirms robust budding of VP40 into the extracellular solution (Fig 3). With WT serving as a phenotypic baseline, we then imaged mutants that abrogate the three main molecular determinants of matrix assembly: the membrane-binding basic patch; the dimerization interface; and the C-terminal oligomerization interface.

**Figure 2:**
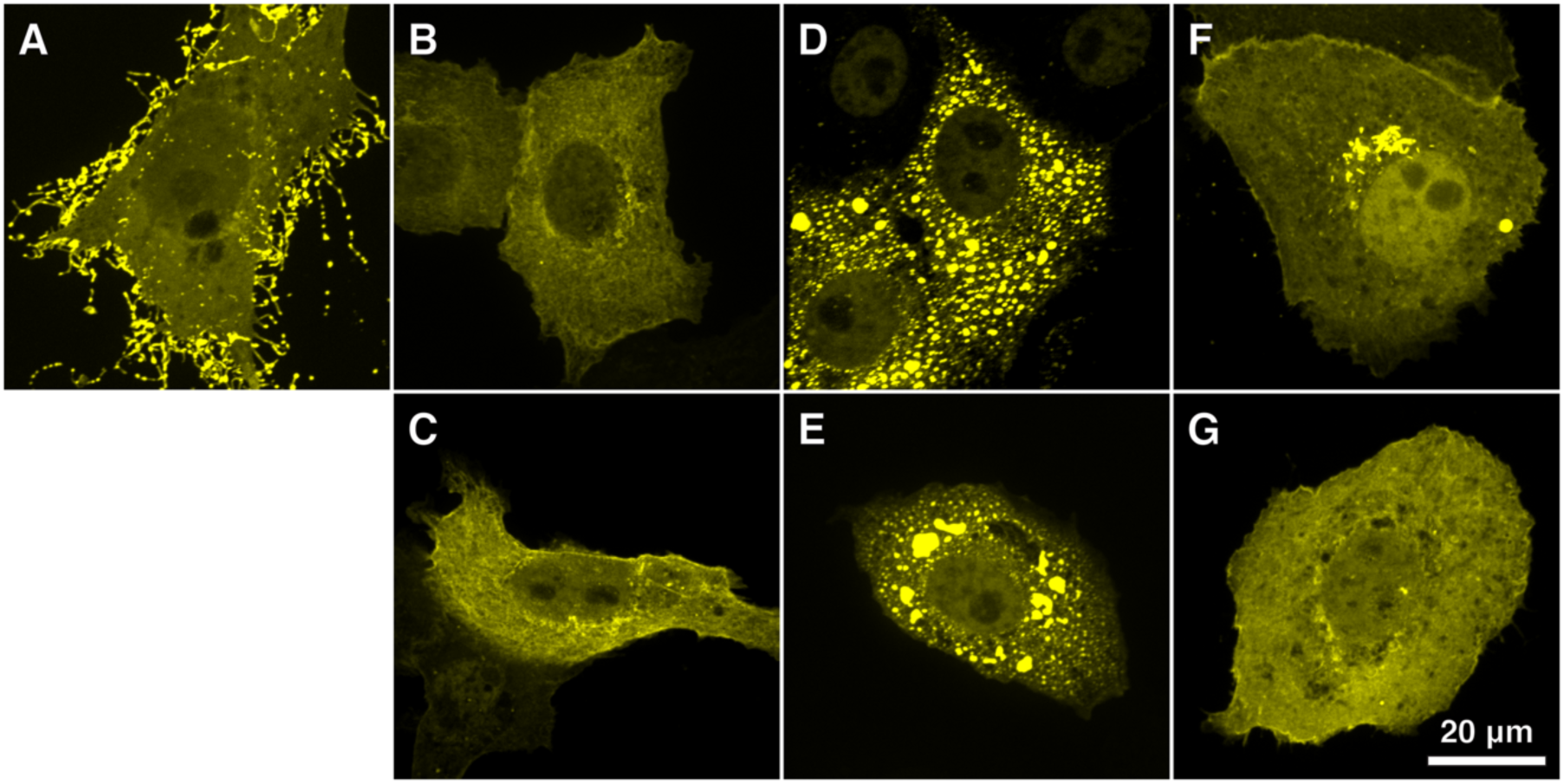
Confocal microscopy of VP40 mutations at the assembly interfaces. A) WT VP40, B) K224/225E, C) K274/275E, D) L117R, E) L117R-K224/225E, F) M241R, G) M241R-K224/225E. Images are maximum-intensity projections with yellow signal indicating sfGFP-VP40 signal.

**Figure 3:**
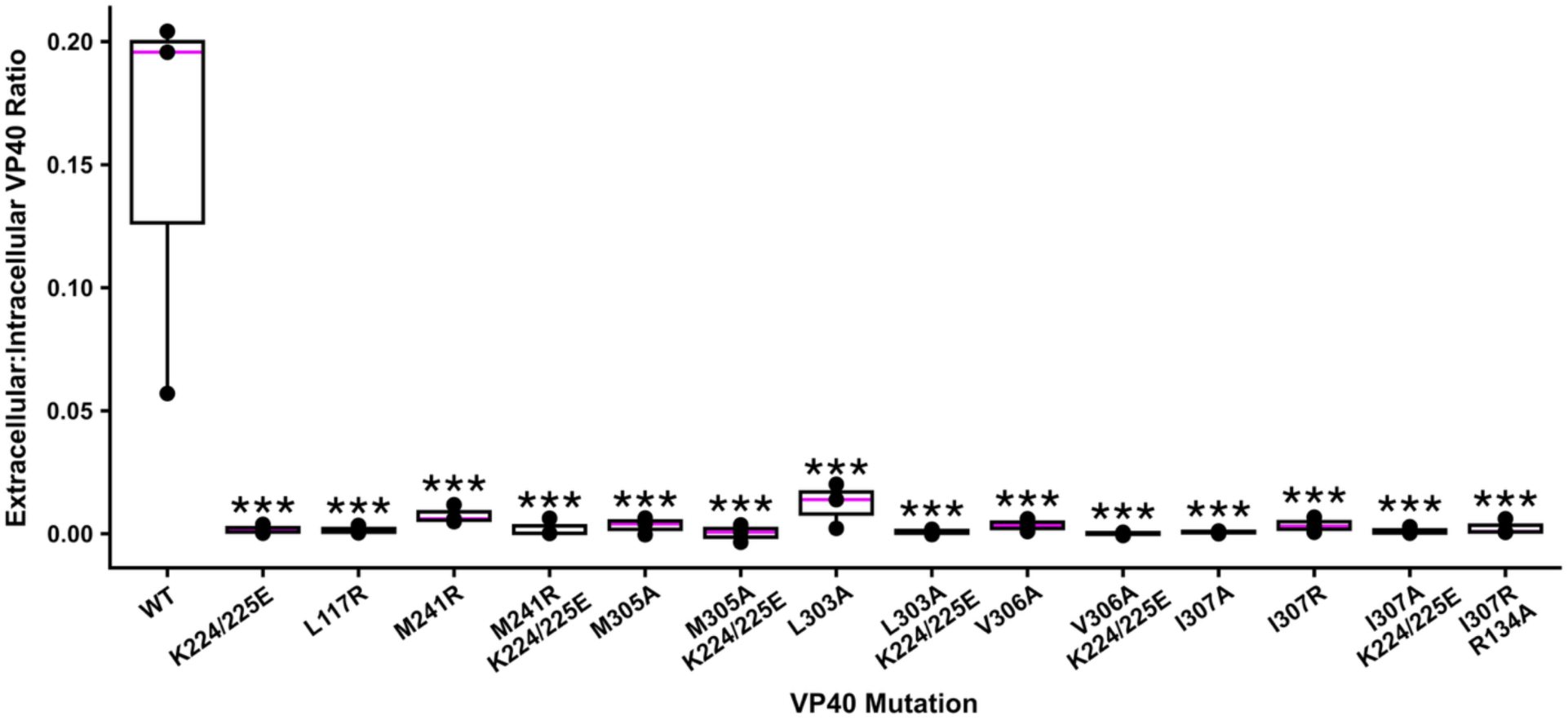
Quantification of Western Blots Showing the Ratio of Extracellular and Intracellular VP40. Western blots were generated as biological triplicates and the ratio of extracellular to intracellular VP40 was quantified for each replicate. Plots are box and whisker plots, with centers showing the median, whiskers showing the range, and box edges indicated the quantiles. *** indicates a P<0.001 diWerence of each mutant with respect to the WT.

For the basic patch, two pairs of mutations were previously described to abrogate membrane binding and budding: mutations from lysine to glutamate at residues 224 and 225 (K224/225E) and residues 274 and 275 (K274/275E) (Fig 1C) (7). In our imaging, each of these mutants produces diWuse signal throughout the cytosol (Fig 2B,C) with no distinct subcellular localization or filament budding. Given the similarity of these two mutations, we decided to use K224/225E as our main membrane-binding mutation. Western blot analysis of K224/225E (Fig 3) also shows minimal VP40 presence in the extracellular solution, demonstrating severely impaired budding.

The L117R mutation is located at the NTD-NTD interface involved in VP40 heterodimer formation (Fig 1C) and was previously shown to prevent dimerization and downstream filament assembly (7). In our images, the L117R mutant shows no filament formation but results in aggregates of various sizes distributed across the cytosol (Fig 2D). Western blot analysis shows severely impaired budding eWiciency of this mutant (Fig 3). Co-mutation of L117R with K224/225E does not aWect the aggregation phenotype (Fig 2D, E), suggesting that these aggregates form freely within the cytosol.

M241R is a mutation located at the C-terminal oligomerization domain (Fig 1C) and was described to preventing viral budding (7). Ultrastructural examination at the cell periphery using resin-embedded electron microscopy suggested that the M241R mutant causes a membrane ruWling morphology without the formation of filamentous buds (7). In our sfGFP-labelled construct, we find that while the M241R mutant does not assemble filaments (Fig 2F), it does generate a distinct aggregation phenotype localized near the nucleus. Western blotting confirms M241R’s impaired budding eWiciency (Fig 3). With the addition of the K224/225E mutation that abrogates membrane-binding, VP40 M241R-K224/225E no longer has any distinct aggregation phenotype (Fig 2G) and instead exhibits the same diWuse signal as the K224/225E mutation alone (Fig 2B), demonstrating that membrane binding is required for this perinuclear aggregation phenotype. Co-expression of M241R with fluorescently-labelled giantin (Fig 4) (20, 21), a cis-Golgi marker, shows spatial colocalization. This along with the loss of subcellular localization with the K224/225E co-mutation suggests that M241R aggregates on the surface of cis-Golgi membranes.

**Figure 4:**
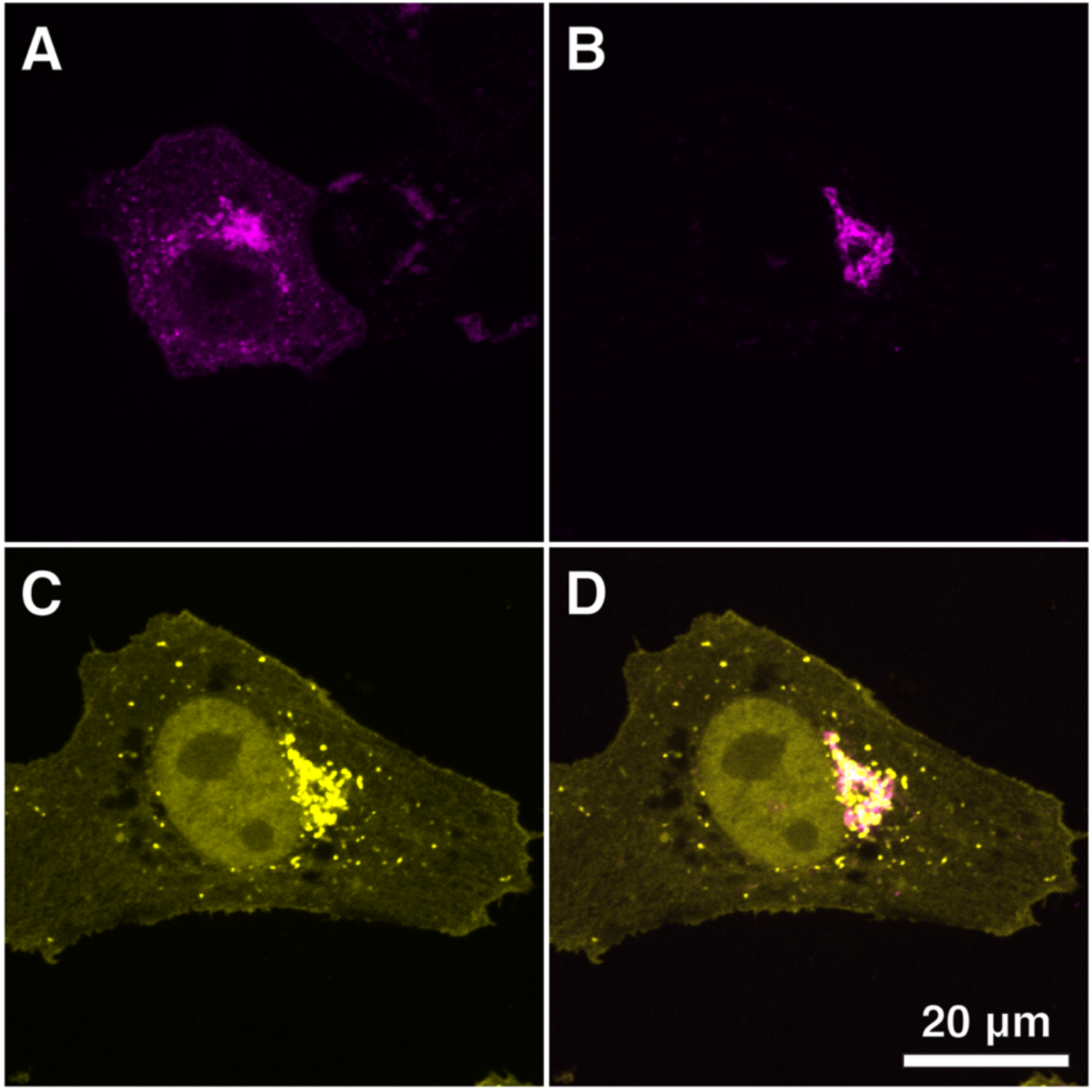
Colocalization of the VP40-M241R Mutation to the Cis-Golgi. A) mScarlet3-giantin expressed alone. B – C) mScarlet3-giantin co-expressed with sfGFP-labelled VP40-M241R. B) mScarlet3-giantin, C) sfGFP-labelled VP40-M241R, and D) overlay of both. Images are of maximum intensity projections of confocal stacks, with yellow indicating sfGFP-VP40, and magenta indicating mScarlet3-giantin.

### Mutations related to Sec24c binding

It was previously shown that VP40 binds directly to Sec24c, a component of the COPII anterograde transport machinery that mediates transport between the ER and Golgi. Knockdown of Sec24c results in reduced budding of VP40 VLPs, suggesting that the VP40-Sec24c interaction is necessary for downstream matrix assembly and budding (16). Scanning alanine mutagenesis isolated residues 303 – 307 as a key area for Sec24c binding. Specifically, co-immunoprecipitation experiments showed that M305A has increased Sec24c aWinity, while mutants L303A, V306A, and I307A have decreased aWinity to Sec24c; T308A showed no eWect.

Imaging of sfGFP-labelled M305A shows that it aggregates close to the nucleus and takes on a ribbon-like phenotype (Fig 5A) that is distinctly diWerent from the perinuclear phenotype of the M241R mutation (Fig 2F). However, like the M241R mutation, M305A shows no signs of filament formation at the cell edges, Western blot analysis shows that M305A produces minimal extracellular VP40 (Fig 3), demonstrating deficiency in budding that is potentially related to inability to traWic to the plasma membrane. With the addition of the K224/225E mutation, M305A-K224/225E no longer has any preferential subcellular localization (Fig 5B) and instead shows the same diWuse cytosolic signal as the K224/225E mutation alone (Fig 2B), indicating that the perinuclear aggregation of M305A requires membrane binding.

**Figure 5:**
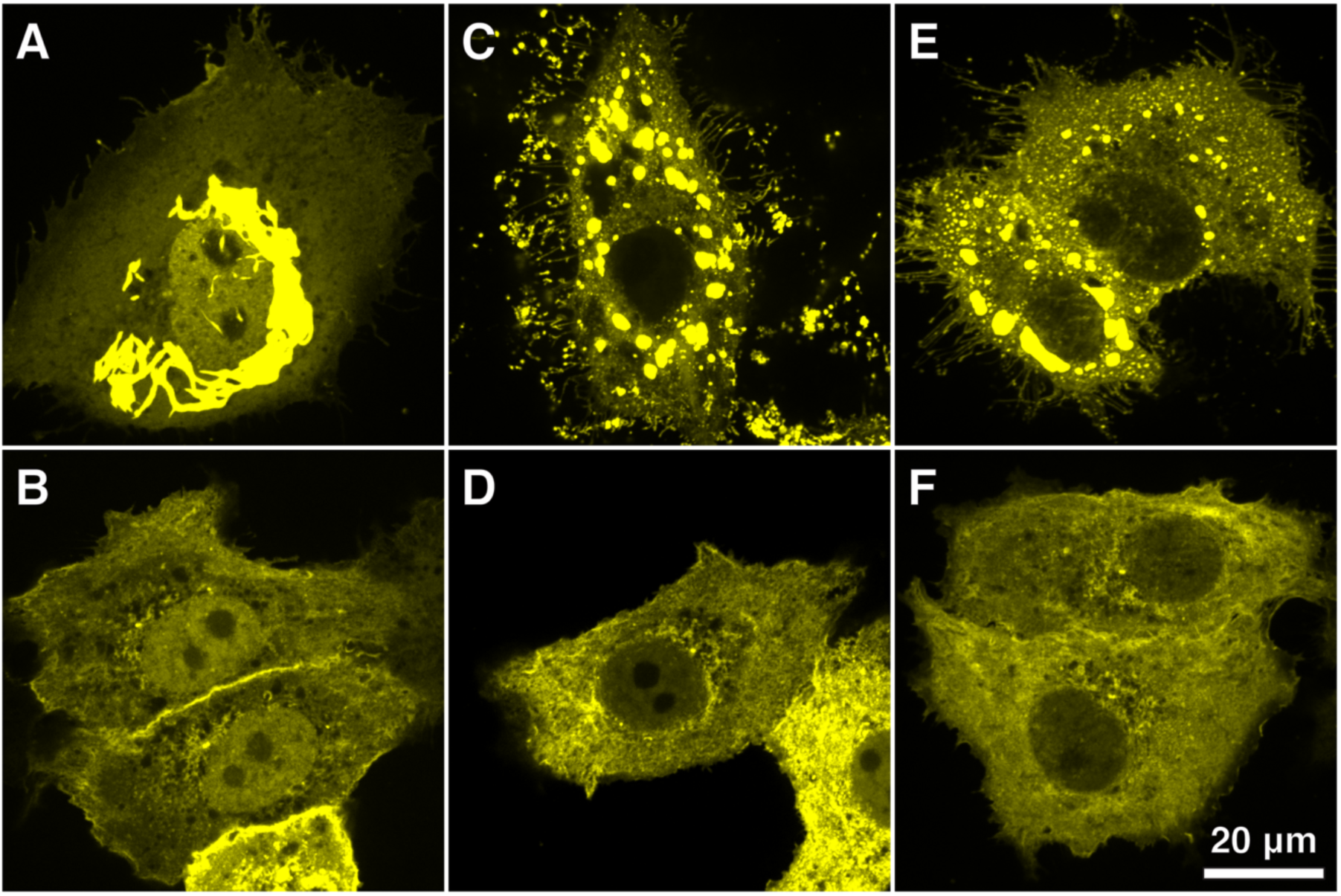
Confocal Microscopy of VP40 Mutations Related to Sec24c Binding. A) M305A, B) M305A-K224/225E, C) L303A, D) L303A-K224/225E, E) V306A, F) V306A-K224/225E. Images are maximum-intensity projections with yellow signal indicating sfGFP-VP40 signal.

Our imaging of L303A and V306A show that they exhibit similar phenotypes, consisting of a combination of large and small puncta distributed in the cytosol as well as filamentous protrusions at the cell edges (Fig 5C,E). While western blot analysis shows that the eWiciency of budding is minimal (Fig 3), the presence of filaments suggests that the lack of budding may not be primarily due to a deficiency in matrix assembly but may instead be due to ineWicient traWicking of VP40 to the PM. Co-mutation of L303A and V306A mutants with K224/225E (Fig 5D,F) results in the loss of aggregates and produces diWuse cytosolic signal consistent with the K224/225E (Fig 2B) mutation alone, showing that the cytosolic aggregation requires membrane binding.

To determine the subcellular localization of L303A, M305A, and V306A, we co-expressed these, along with WT, with diWerent fluorescently-labelled subcellular markers.

For the ER, we used mCherry-labelled calnexin (Fig 6). When expressed alone, the calnexin surrounds the nucleus and shows relatively-dense signal that terminates well before the cell edges. Co-expression with WT shows a similar calnexin distribution but also contains weaker signal that extends towards the PM. Co-expression of calnexin with L303A and V306A show similar phenotypes with WT, with no colocalization with the subcellular aggregates. While M305A shows no clear colocalization, the distribution of calnexin is notably diWerent from that with WT co-expression. M305A expression causes calnexin signal to spread out towards the PM and also within the nucleus, suggesting that M305A causes disruption to the ER.

**Figure 6:**
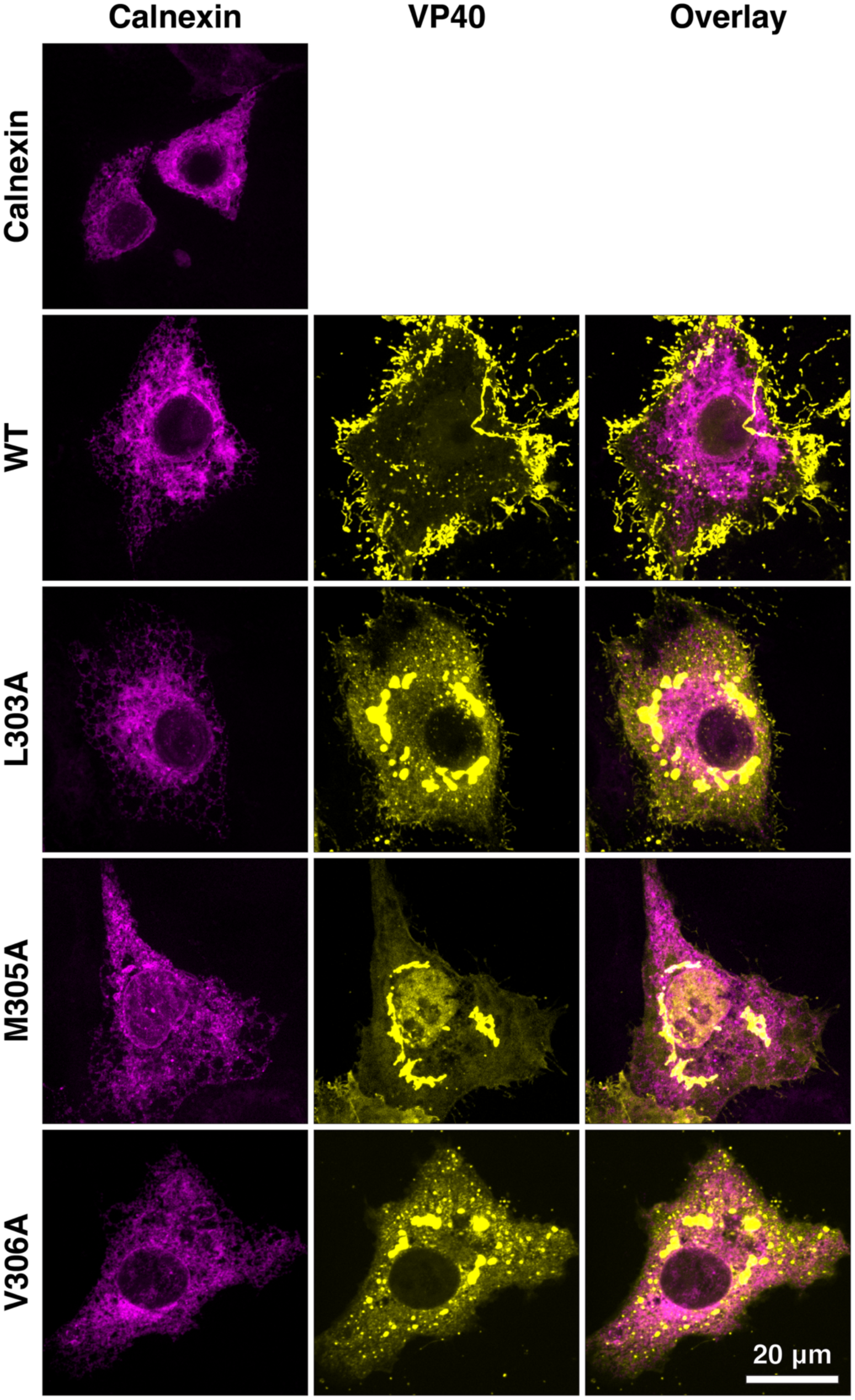
Co-expression of calnexin with Sec24c-related VP40 mutants. Images are maximum-intensity projections with yellow signal indicating sfGFP-VP40 signal and magenta signal indicating mCherry-calnexin.

For the endoplasmic-reticulum–Golgi intermediate compartment (ERGIC), we used mScarlet3-ERGIC53 as a marker (Fig 7). When expressed alone, ERGIC53 primarily localizes in a patch near the nucleus with some puncta throughout the cytosol. Co-expression with WT, L303A, M305A, and V306A do not show any significant changes in ERGIC53 localization, suggesting that the aggregates formed by these VP40 mutants occur outside of the ERGIC and do not significantly interact with ERGIC proteins.

**Figure 7:**
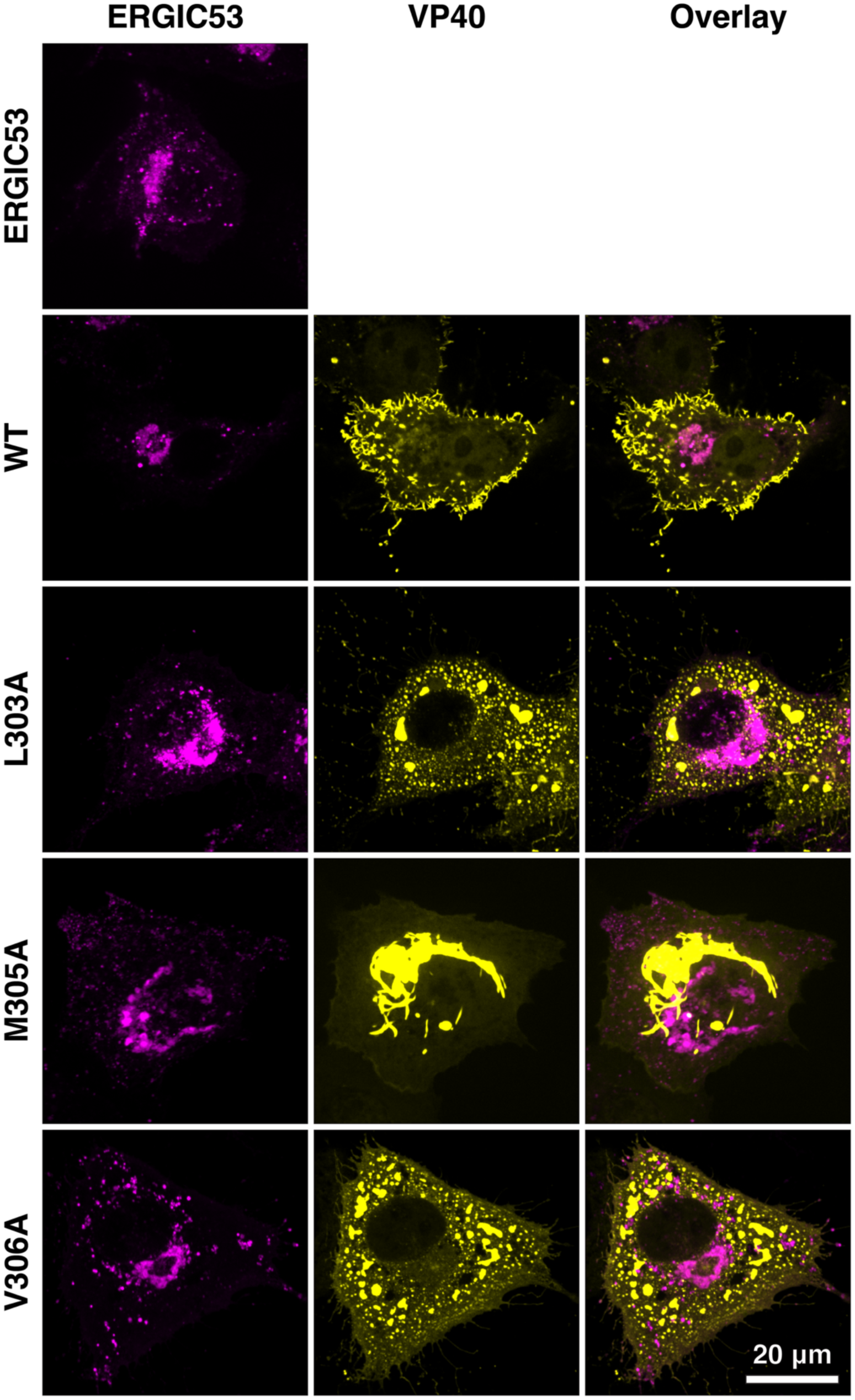
Co-expression of ERGIC53 with Sec24c-related VP40 mutants. Images are maximum-intensity projections with yellow signal indicating sfGFP-VP40 signal and magenta signal indicating mCherry-ERGIC53.

For the cis-Golgi, we used mScarlet3-giantin as a marker (Fig 8). As described above for M241R, giantin primarily localizes near the nucleus with some cytosolic signal. WT, L303A, M305A, and V306A do not show any direct colocalization similar to M241R, suggesting that none of these mutants aggregate directly on the cis-Golgi. However, M305A does seem to cause increased giantin signal across the cytosol, suggesting that M305A may produce some low level of cis-Golgi disruption.

**Figure 8:**
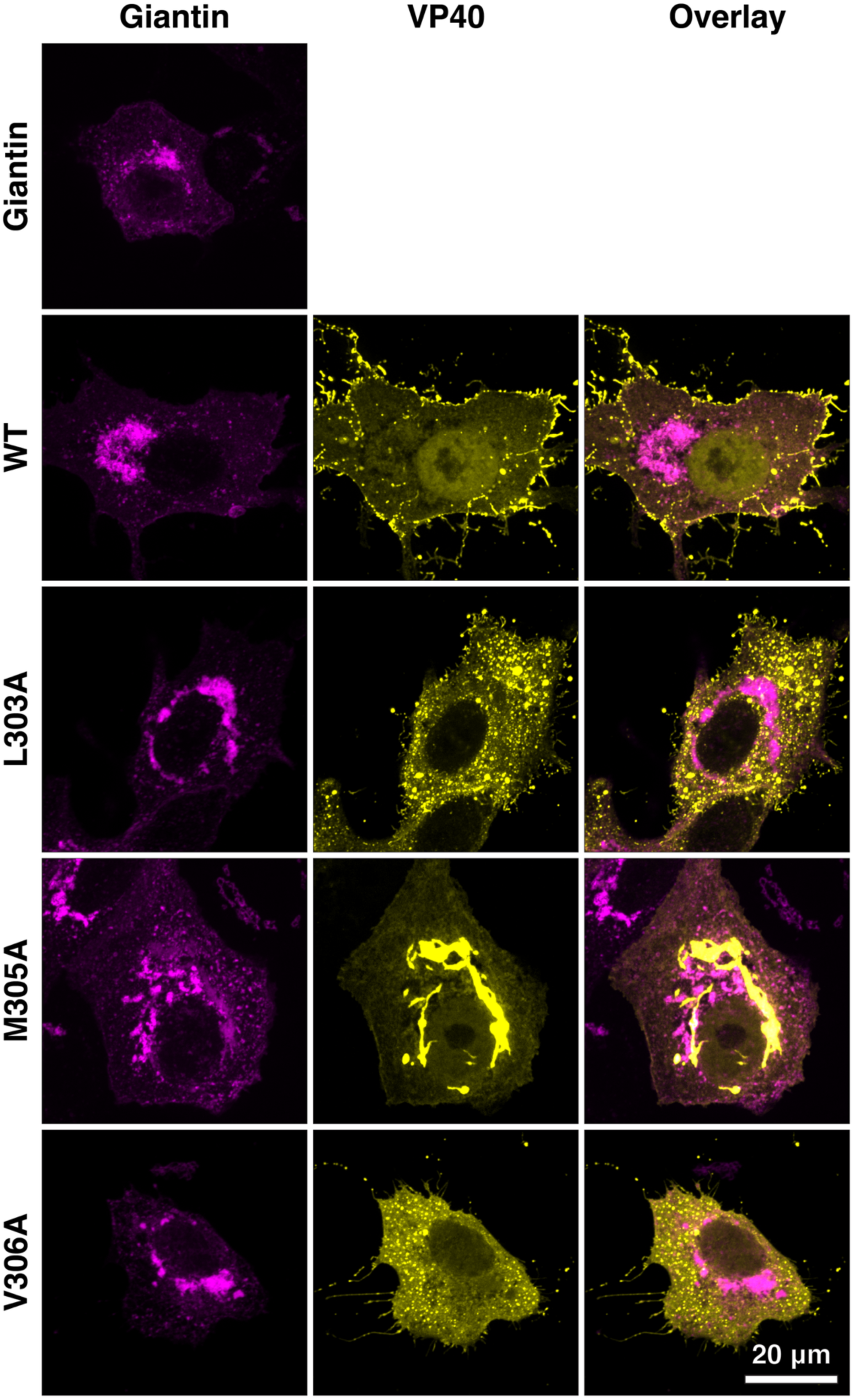
Co-expression of giantin with Sec24c-related VP40 mutants. Images are maximum-intensity projections with yellow signal indicating sfGFP-VP40 signal and magenta signal indicating mScarlet3-giantin.

For the trans-Golgi network, we used mCherry-labelled trans-Golgi network protein 2 (TGNP, also known as TGOLN2) (Fig 9). TGNP alone shows perinuclear signal as well as puncta distributed throughout the cytosol. Co-expression with WT and M305A show TGNP distribution similar to when it is expressed alone. While co-expression of L303A and V306A also shows TGNP signal in the perinuclear and cytosolic regions, similar to when TGNP is expressed alone, L303A and V306A cause a substantial increase in nuclear-localized TGNP. Interestingly, the nuclear localization of L303A and V306A is reduced compared to WT and M305A, suggesting that they do not induce the nuclear localization of TGNP by direct binding of mutant VP40 but instead by disrupting the trans-Golgi network. WT shows low levels of nuclear TGNP signal as compared to L303A and V306A, while M305A does not seem to have any nuclear TGNP signal.

**Figure 9:**
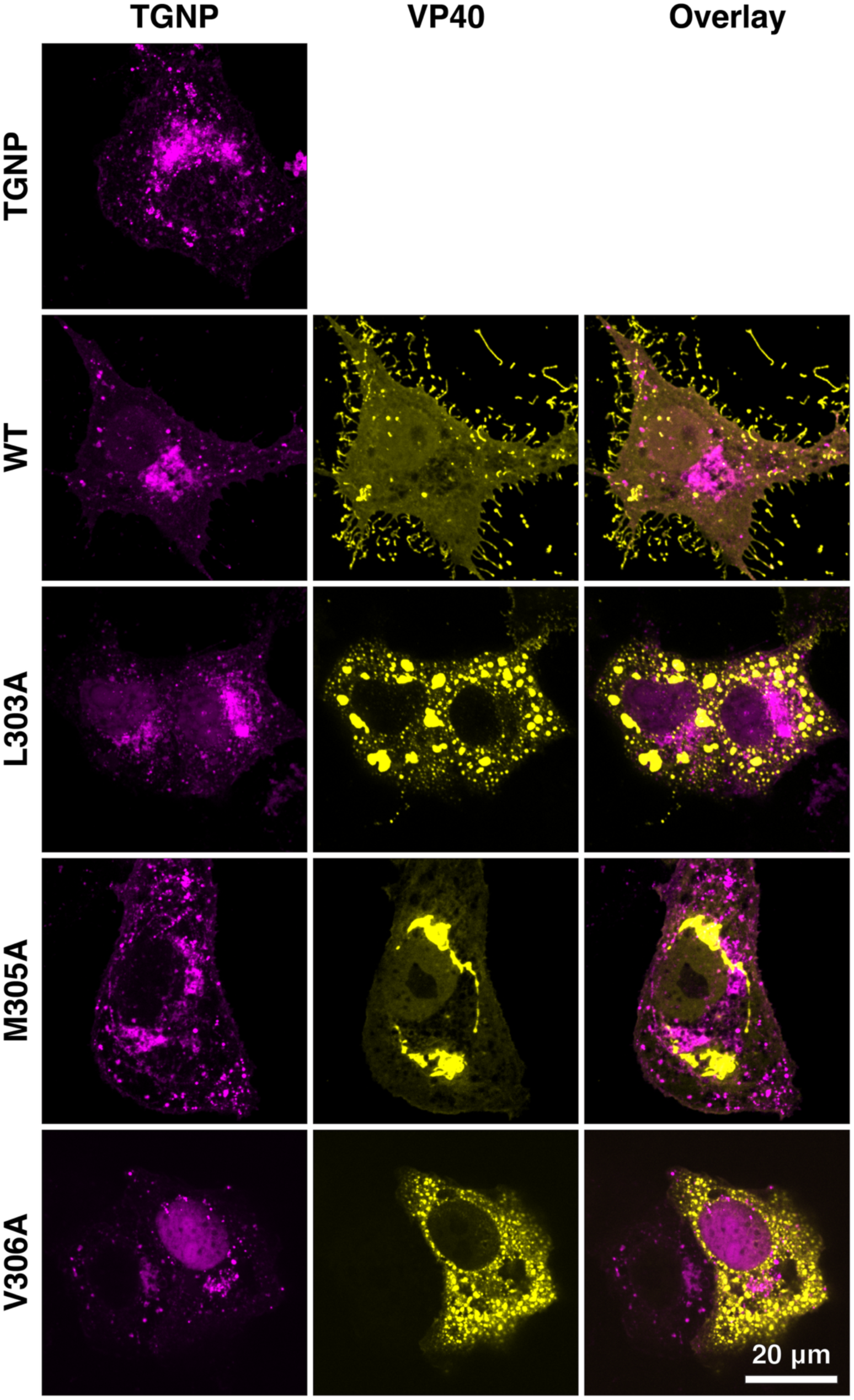
Co-expression of TGNP with Sec24c-related VP40 mutants. Images are maximum-intensity projections with yellow signal indicating sfGFP-VP40 signal and magenta signal indicating mCherry-TGNP.

### Mutations to I307

While the I307A mutation was described as being Sec24c binding deficient (16), a similar I307R mutation was described as causing VP40 to prefer the RNA-binding octameric conformation (7). Our imaging of both these mutants shows a consistent phenotype between the two consisting of a combination of large and small cytosolic aggregates (Fig 10A,B). Addition of the K224/225E mutation to I307A shows the same pattern of cytosolic aggregation (Fig 10C), indicating that unlike the other Sec24c mutants described above, the aggregation of this mutant is not dependent on the ability to bind membranes. An additional R134A mutation was previously shown to abolish RNA-binding of the I307R mutation (7). In our results, we find that adding the R134A co-mutation to I307R reduces the presence of small cytosolic puncta but not the large ones (Fig 10D). Western blot analysis of these mutants show minimal VP40 budding (Fig 3). Taking both the K224/225E and R134A co-mutations into account, our results suggest that the large scale aggregation of I307 mutants is unrelated to RNA or membrane binding and instead represent some other form of cytosolic aggregation. The phenotype of these aggregates resemble the phenotypes of the L117R and L117R-K224/225E mutants (Fig 2D,E), suggesting they may be similar in nature.

**Figure 10:**
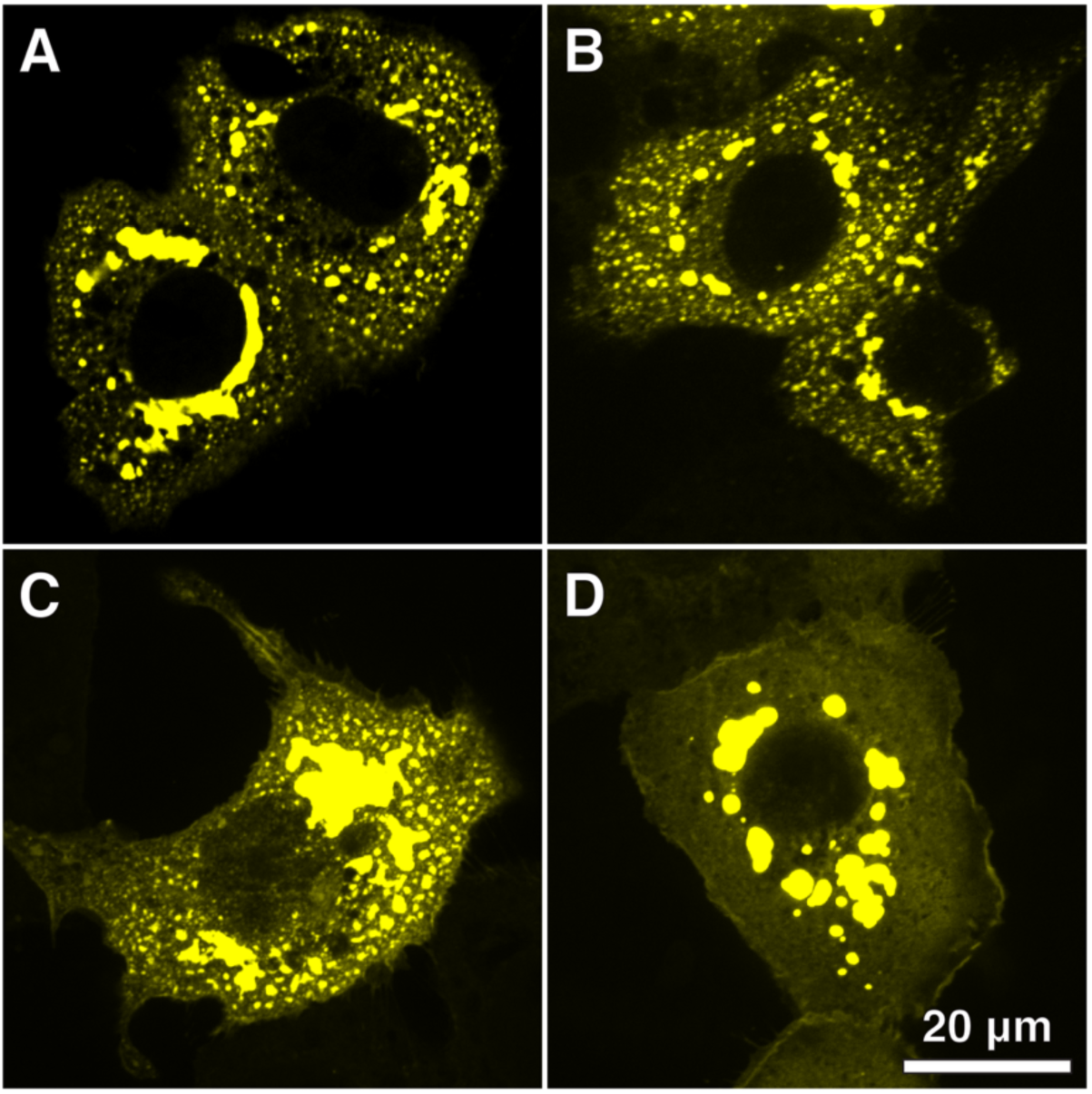
Confocal Microscopy of VP40 Mutations to I307. A) I307A, B) I307R, C) I307A-K224/225E, D) I307R-R134A. Images are maximum-intensity projections with yellow signal indicating sfGFP-VP40 signal.

## Discussion

A key function of VP40 in the Ebola virus life cycle is to serve as the viral matrix protein, which plays an essential role in the budding of Ebola’s characteristic filamentous particles (5). VP40 binds the host PM and its assembly into the 2D-crystalline matrix layer causes the PM to curve into filamentous buds (4). During matrix assembly, VP40 packages nucleocapsids or nucleocapsid-like assemblies in budded viral particles (22, 23), but in the absence of nucleocapsids, VP40 readily buds empty VLPs (4, 5). This demonstrates that VP40 alone is suWicient to drive all necessary VP40-host interactions required for matrix assembly and budding. While VP40 is expressed throughout the host cytosol, it does not appear to directly bind the PM and instead seems to require host traWicking machinery to arrive at the PM. In this manuscript, we have used sfGFP-labelled VP40 mutants to visualize how mutations aWect VP40’s subcellular localization, which has provided new insights into how VP40 traWics from the cytosol to the PM.

Our sfGFP-labelling system provides improved resolution of subcellular localization over immunolabelling approaches, as it does not require the use of detergents for membrane permeabilization. This seems to be a an important factor, as many of the mutants we characterized are membrane-tethered and permeabilization may disrupt the subcellular phenotypes of these mutants. A prime example of this is the M241R mutant, which was previously characterized by resin-embedded electron microscopy as leading to ruWling at the PM rather than filament formation, suggesting that impaired budding is due only to a deficiency in matrix assembly (7). While M241R likely does impair matrix assembly, our confocal imaging showed that M241R aggregates near the nucleus and co-expression with labelled giantin indicated that M241R localizes at the cis-Golgi. Co-mutation of M241R with K224/225E results in diWuse cytosolic signal, showing that aggregation is dependent on membrane binding. These observations suggest that aggregation and impaired traWicking to the PM are also likely major factors in the deficient budding of the M241R mutation; such observations required using imaging approaches that visualize the distribution of VP40 throughout the whole cell.

We also characterized a set of mutations spanning residues 303 – 307, which were shown to modify binding aWinity with Sec24c using biochemical methods (16). Sec24c is a key component of COPII coats, which are involved in anterograde traWicking between the ER and Golgi. Genetically-encoded labels and confocal microscopy allow for whole-cell imaging, providing a more holistic view on the cellular phenotypes of these VP40 mutants beyond protein-protein interactions. We find that M305A, which was biochemically shown to have increased binding aWinity to Sec24c, aggregates with a distinct ribbon-like morphology near the nucleus and disrupts the subcellular localization of calnexin, an ER marker. The L303A and V306A mutants, which have been shown to have decreased Sec24c aWinity, form punctate aggregates distributed throughout the cytosol and cause mis-localization of TGNP, a trans-Golgi network marker, to the nucleus. Co-mutation of these three mutants with K224/225E results in cytosolic distribution, showing that their aggregation only occurs when VP40 is capable of binding membranes. Taking account this need for membrane binding, along with our observations on M241R, we believe that the phenotypes of these mutants represent stalled traWicking intermediates.

We identified two I307 mutants in the literature, with I307A characterized as reducing aWinity to Sec24c (16) and I307R characterized as preferring VP40’s RNA-binding octameric isoform (7). Our imaging showed that these two mutants have indistinguishable subcellular phenotypes consisting of small and large punctate aggregates throughout the cytosol. Co-mutation of I307A with K224/225E showed that these aggregates do not require membrane binding and co-mutation of I307R with R134A, which prevents RNA-binding, only results in the loss of small puncta. This suggests that mutations to I307 seem to cause more general cytosolic aggregation unrelated to Sec24c- or RNA-binding, and likely do not represent a stalled traWicking intermediate. The I307A and I307R phenotypes are also similar to the L117R mutant, which disrupts the formation of VP40 heterodimers, which may suggest a similar mechanism for aggregation.

Despite eWorts to determine the localization of our VP40 mutants by co-expressing localization markers, only M241R showed distinct colocalization with a marker. This may be due to substantial changes in membrane architectures or the occlusion of other membrane-associated proteins due to the high level of VP40 aggregation. WT VP40 does not exhibit any discernable subcellular localization, suggesting that traWicking of VP40 typically occurs at low local concentrations that are not discernable above free-floating cytosolic VP40. For future studies, direct molecular-resolution imaging by cryo-ET may be important for determining the subcellular localization of VP40 mutants at expression levels where minimal aggregation occurs.

Given the necessity of Sec24c binding for budding and the subcellular localization of our mutants, we suggest a possible mechanism of VP40 traWicking from the cytosol to the PM. M305A’s enhanced aWinity to Sec24c, aggregation localized around the nucleus, and disruption of calnexin signal suggests that it may be stalled at the ER, at the origin of COPII coat formation and beginning of its traWicking pathway. M241R potentially modulates some other VP40 host-protein interaction, resulting in it being stalled further in the secretory pathway at the cis-Golgi. The more disperse cytosolic distribution of L303A and V306A suggest stalling further along VP40’s traWicking pathway; the mis-localization of TGNP caused by these mutants suggest that they impair the trans-Golgi network. Throughout this process, VP40 needs to be membrane-tethered for traWicking. Initial binding to Sec24c at the ER may bring the membrane-binding polar regions of VP40 into close apposition to the ER membrane, facilitating the initial membrane tethering event. As VP40 traWics, its relative aWinity to diWerent unknown traWicking proteins allows it to progress past the anterograde transport of COPII vesicles, which terminates at the cis-Golgi, eventually allowing VP40 to arrive that the PM. Given the cytosolic expression of VP40, this also suggests that VP40 is traWicked along the outside of vesicles; fusion of such vesicles at the PM would result in VP40 tethered to the inner leaflet, where it would then assemble into the matrix layer. Recent structural work on COPII vesicles suggests that they are only sparsely coated (24), which may provide suWicient areas of uncoated membrane to traWic small amounts of VP40.

## Acknowledgements

This work was supported by U.S. National Institutes of Health grant DP2GM146321 (to WW). WW is also a Pew Scholar in the Biomedical Sciences and supported by the Pew Charitable Trusts. TH was supported by the U.S. National Institutes of Health Molecular Biophysics Training Grant at Vanderbilt T32GM008320. Confocal microscopy was performed in the Vanderbilt Cell Imaging Shared Resource (supported by NIH grants CA68485, DK20593, DK58404, DK59637 and EY08126).

## Materials and Methods

### Plasmids, Cloning, and Site-Directed Mutagenesis

VP40 used in this study is from the Zaire Ebolavirus Mayinga-76 strain. It is N-terminally labelled using sfGFP (25) and was cloned into a pCDNA5/TO vector (Thermo Fisher Scientific) using Gibson assembly (26). To generate VP40 mutants, site-directed mutagenesis was performed using the New England Biolabs Q5 strategy (New England Biolabs) with primers generated by NEBaseChanger. All constructs were validated using whole-plasmid Oxford nanopore sequencing (Plasmidsaurus).

The mScarlet-Giantin construct used was Addgene plasmid #85048. The mCherry-calnexin construct was Addgene plasmid #55005. The mScarlet-ERGIC53 construct was cloned using an mEmerald-ERGIC53 construct (Addgene #170717) as the base vector, with the mEmerald swapped for mScarlet via Gibson cloning. The mCherry-TGNP construct was Addgene plasmid #55145.

### Cell Culture, Transfection, and Preparation for Confocal Microscopy

Vero E6 cells were maintained in Minimal Essential Medium (MEM) supplemented with 10% fetal bovine serum (FBS) and grown at 37°C with 5% CO_2_. For confocal imaging, Vero E6 cells were seeded in glass bottom 12-well plates (Cellvis P12-1.5H-N) at approximately 70,000 cells / well to obtain 60-80% confluency 24-hours post seeding. Plasmids were transfected into Vero-E6 cells using Mirus TransIT transfection reagent (Mirus). Cells were transfected for 18 hours prior to fixation with 4% paraformaldehyde (PFA) in phosphate-buWered saline (PBS) for 15 minutes at room temperature. After fixation, cell nuclei were stained with NucBlue Fixed Cell ReadyProbes Reagent (Thermo Fisher Scientific) containing 4′,6-diamidino-2-phenylindole (DAPI) for 5 minutes at room temperature. Fixed cells were stored at 4°C in PBS and covered from light prior to imaging.

### Spinning Disc Confocal Microscopy

Images were collected at the Vanderbilt Nikon Center for Excellence using the Nikon Spinning Disc confocal microscope provided by the Vanderbilt Cell Imaging Shared Resource (CISR) core. An Andor DU-897 EMCCD camera was used to capture images at 100x through an Apo TIRF (oil) 100x 1.49 NA WD 0.12mm objective. All images were collected as Z-stacks and displayed as maximum intensity projections.

### Western Blotting and Quantitation

All steps were performed as biological triplicates. Vero E6 cells were transfected using Mirus TransIT transfection reagent (Mirus) using a reverse transfection approach (27) and seeded into plastic 6-well plates (Sarstedt) at a concentration of 200,000 cells / well. Cells and supernatant were harvested 48 hours post transfection.

For extracellular VP40, cell supernatants were collected and mixed with Halt Protease Inhibitor Cocktail (Thermo Fisher Scientific) and 5 mM Ethylenediaminetetraacetic acid (EDTA) and stored on ice. Supernatant was then clarified by centrifugation at 800×g for 5 minutes at 4°C. Clarified supernatants were then added to a 20% (w/v) sucrose cushion in Tris-buWered saline (TBS) and ultracentrifuged at 110,000×g for 1.5 hours at 4°C using a SW41Ti rotor and Optima XE-90 Ultracentrifuge (Beckman Coulter). The supernatant and cushion were removed and the pellet was resuspended in 50 mM tris(2-carboxyethyl)phosphine (TCEP), 4M urea, 1.8× LDS gel-loading buWer (Invitrogen) and heated in a 95°C heat block for 5 minutes.

For intracellular VP40, cells in each well were washed with PBS and incubated with 0.25× Trypsin-EDTA at 37°C for 20 minutes. Trypsin-EDTA was neutralized using MEM supplemented with 10% FBS and cells were collected and centrifuged at 200×g for 5 minutes at 4°C. To wash oW any residual trypsin, cell pellets were resuspended in PBS and centrifuged again at 200×g for 5 minutes at 4°C. Cell pellets were then lysed by resuspension in radioimmunoprecipitation assay (RIPA) buWer (Sigma-Aldrich) and Halt Protease Inhibitor Cocktail and EDTA and incubated on ice for 5 minutes. Cell debris was removed by centrifugation at 13.3×g for 10 minutes at 4°C.

To normalize for cellular protein content, bicinchoninic acid assay (BCA) assays (Thermo Fisher Scien5fic) were performed on cell lysates in a 96 well plate following manufacturer’s protocols; plates were read using a Molecular Devices SpectraMax iD3 plate reader (Molecular Devices). Samples for each mutant were normalized to the lowest protein concentra5on in the replicate and gel samples were prepared with 50 mM TCEP, 1× LDS buffer in PBS, incubated at room temperature for 5 minutes and then heated at 95°C for 5 minutes. Extracellular samples were diluted to match normaliza5on factor of the lysate samples.

Samples were run loaded and ran on NuPAGE 4-12% Bis-Tris gels (Invitrogen) for 1.75 hours on ice at 100V. Gels were semi-dry transferred to nitrocellulose membranes using a Trans-Blot Turbo Transfer System (Bio-Rad) at 25V for 10 minutes. Membranes then were blocked using filtered 5% BSA in TBS blocking buWer for 1 hour at room temperature while being rocked. Then, membranes were incubated in mouse anti-Ebola Virus VP40 monoclonal antibody (Invitrogen MA5-47499) diluted 1:1000 in 5% BSA in TBS overnight at 4°C while rocking. Membranes were washed 3 times in TBS + 0.1% Tween-20 (TBST) for 5, 10, and 15 minutes and then washed in TBS for 5 minutes while rocking at room temperature.

Membranes were incubated in goat anti-mouse IRDye 680LT antibody (LI-COR) diluted 1:10,000 in 5% BSA in TBS for 1 hour at room temperature while rocking. Membranes were washed 3 times in TBST for 5, 10, and 15 minutes and then washed in TBS for 5 minutes while rocking at room temperature. Western blots were imaged using LI-COR Odyssey CLx at 700 nm (LI-COR). Blots were in quantified in ImageJ (28), using the BandPeakQuani5fica5on macro (29) with parameters of 1 pixel background width, top/boZom and sides used for es5ma5ng background, and background es5ma5on by mean. In The ra5o of detected VP40 in the intra- and extracellular samples was calculated in RStudio (30). An ANOVA test was performed (p-value < 0.05) and DunneZ’s test was used as a post-hoc analysis to compare extracellular:intracellular VP40 ra5os of WT VP40 to VP40 mutants.

## Notes

### Competing Interest Statement

The authors have declared no competing interest.

### Summary of Updates

This version has been revised to update the figures from the previous draft as well as to add additional experiments with co-expression of mutant VP40 with subcellular markers.

