## Supplementary Figure 1 for "Subcellular Characterization of the Molecular Determinants of Ebola VP40 Trafficking and Assembly"

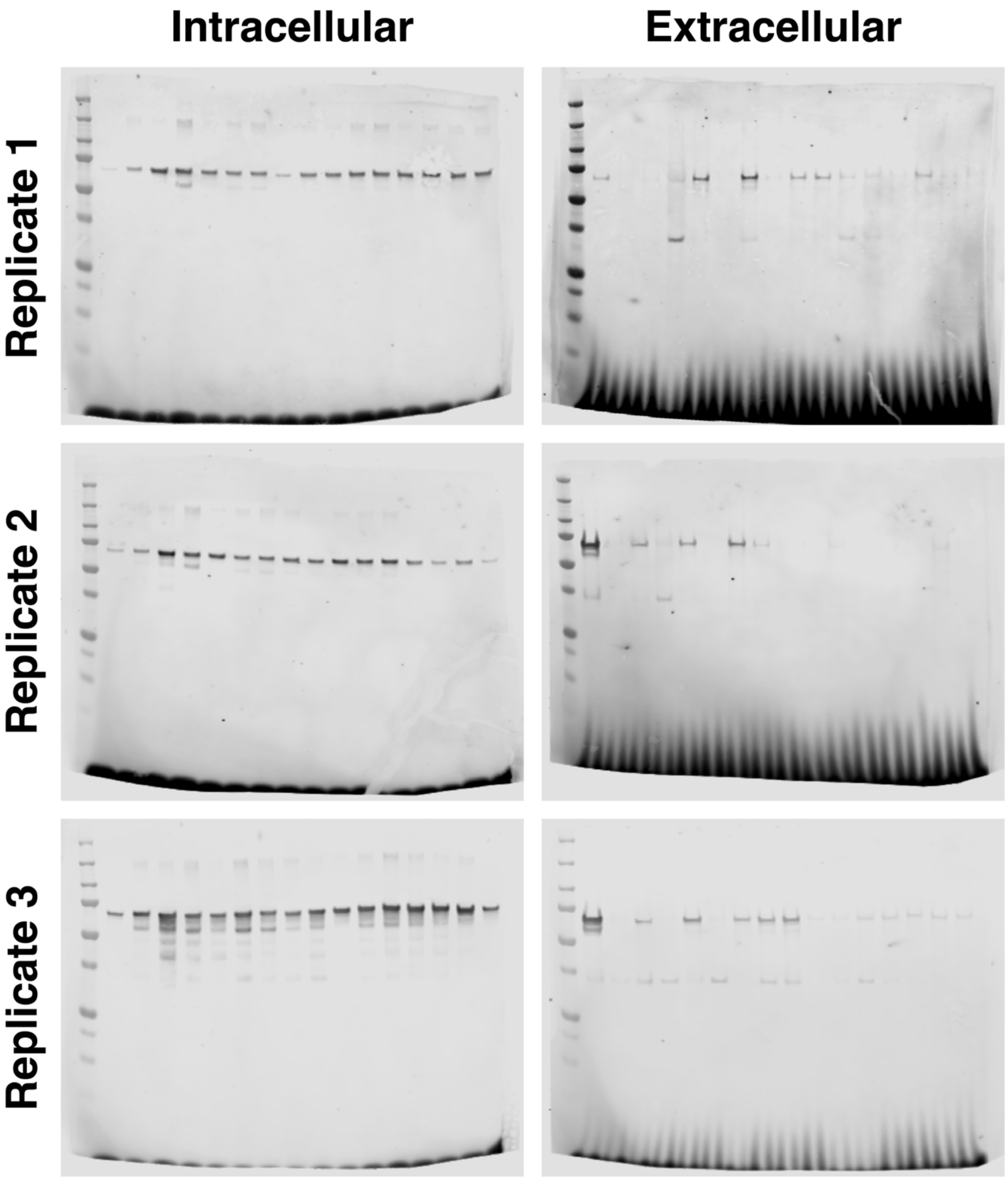


**Supplementary Figure 1: Raw western blot images of intracellular and extracellular VP40.** From left to right, lanes are ladder, WT, L117R, K224/225E, I307R, M241R, I307A, L303A, V306A, M305A, M241R-K224/225E, I307A-K224/225E, L303A-K224/225E, V306A-K224/225E, M305A-K224/225E, I307R- K224/225E, I307A-R134A. Molecular weights of the ladder from top to bottom are 250, 150, 100, 75, 50, 37, 25, 20, 15, 10 kDa.
